# A quasi-paired cohort strategy reveals the impaired detoxifying function of microbes in the gut of autistic children

**DOI:** 10.1101/2020.03.10.984872

**Authors:** Mengxiang Zhang, Yanan Chu, Qingren Meng, Rui Ding, Xing Shi, Zuqun Wang, Yi He, Juan Zhang, Jing Liu, Jie Zhang, Jun Yu, Yu Kang, Juan Wang

**Affiliations:** Department of Biomedical Informatics, School of Basic Medical Sciences, Peking University, Beijing, 100191, China; CAS Key Laboratory of Genome Sciences and Information, Beijing Institute of Genomics, Chinese Academy of Sciences, 100101, Beijing, PR China; Autism research center of Peking University Health Science Center, Beijing, 100191, China; Department of Respiratory & Critical Care Medicine, Peking University People’s Hospital, Beijing, 100044, PR China; Department of pediatrics, Peking University Third Hospital, Beijing, 100191, China; Key Laboratory of Mental Health, Ministry of Health, Peking University Institute of Mental Health, Peking University Sixth Hospital, Beijing, 100191, China; Children health care center, Xi’an Children’ Hospital. Xian, 710003, China

## Abstract

Growing evidence suggests that autism spectrum disorder (ASD) is highly associated with dysbiosis in the gut microbiome. However, results of metagenome-based microbiome studies are not always consistent due to great individual diversity that overwhelms disease-associated alterations. Here, we proposed a novel analysis strategy—quasi-paired cohort and applied it to a metagenomic study of ASD microbiomes. By comparing the paired samples of ASD and neurotypical subjects, we identified significant deficiencies in ASD children in detoxifying enzymes and pathways, which showed strong correlations to mitochondrial damage. Diagnostic models with these detoxifying enzymes accurately discriminated ASD individuals from controls, and the dysfunction score inferred from the model increased with the clinical rating scores of ASD. Conclusively, our findings suggest a previously undiscovered mechanism in which impaired microbial detoxification leads to toxicant accumulation and mitochondrion damage contributes to the pathogenesis of ASD. This novel mechanism points to future therapeutic strategies of rebuilding microbial detoxification for ASD.

## Introduction

Autism spectrum disorder (ASD) is a complex neurodevelopmental disorder, characterized by impaired social communication, and repetitive and stereotyped behavior, interests, or activities(*1*). The prevalence of ASD has been increasing worldwide, and is estimated at one in 59 eight-year-olds in America by the Autism and Developmental Disabilities Monitoring (ADDM) Network in 2014(*2*). The etiology of ASD remains unknown and likely involves a wide range of environmental factors that affect many physiological processes in genetically sensitive individuals(*3*). Recently, lines of evidence showed possible contributions of intestinal microbiomes to the pathogenesis of ASD as in many other diseases(*4*). First, gastrointestinal co-morbidity is common in ASD children(*5*). Second, ASD patients are mostly picky-eaters due to sensory problems or to avoid severe food allergy(*6*) and are often deficient in digestive enzymes(*7*), which unavoidably alters the nutrients or media of the microbes inhabiting their gut and influences the growth of component species, leading to dysbiosis. Finally, many studies have found obvious dysbiosis in the gut microbiome of ASD patients, such as deficiency in *Bifidobacterium longum*, and overgrowth of *Clostridium* spp. and *Candida albicans* (*8, 9*). The dysbiosis of the ASD microbiome is believed to be associated with inflammation in intestinal epithelia and increased permeability of the gut-blood barrier (*10*). However, the mechanisms of how the intestinal microbiome affects ASD pathogenesis are not fully understood.

Metagenome research based on shotgun sequencing is widely applied in exploring microbe-host interactions and microbiome-associated pathogenesis in many diseases. Compared to targeted sequencing of 16S rDNA, which only presents species profiles, shotgun sequencing is more informative as it provides comprehensive information for the inference of concrete metabolic pathways and physiological functions of the microbiome(*11*) (*12*). However, successful shotgun-based metagenome analysis has not been reported in studies of the ASD microbiome. One of the obstacles to metagenomic analysis is the great individual diversity among samples, as the microbiome is influenced by a wide range of factors such as genetics, age, diet, and health, which is hard to control(*13*). The individual diversity is often so great that it even overwhelms disease-associated alterations and highly impacts the identification of disease-associated microbiome features, meaning the results of studies often include stochastic false positives or negatives that are highly dependent on the samples recruited (*14*) (*15*).

It is well known that the microbiome constituents are widely different even among healthy people(*16*). Moreover, microbial components and their abundance are greatly influenced by complex metabolic interactions and strictly constrained by the entire metabolic network of the microbiome(*17, 18*). In this way, relative activity of a specific pathway should be compared between samples of similar metabolic background(*15, 19*). Based on this concept, we developed a novel strategy for metagenomic analysis, i.e. “quasi-paired cohort” where we paired ASD and control samples of similar metabolic background and thus transformed the original group cohort into a paired cohort. In this way, we not only reduced the individual diversity but also increased the statistical power with the same sample size. We then performed shotgun-based metagenome sequencing of 79 fecal samples from ASD individuals and healthy controls, and with the new strategy, we were able to identify apparent deficiencies in pathways of deoxidation and toxicant-degradation in ASD microbiomes. As toxin exposure has been epidemiologically demonstrated as one the major etiological factors of ASD(*20*), impairment of detoxification function of the intestinal microbiome provides a previously unpublished mechanism for why ASD patients are more vulnerable to toxicant exposure and how the intestinal microbiome contributes to the pathogenesis of ASD.

## Materials and Methods

### Study Design

ASD children and gender- and age-matched neurotypical controls were consecutively recruited from the Autism Research Center of Peking University Health Science Center and surrounding communities. Fecal and morning urine samples were collected from each participant for metagenome sequencing and urine metabolite measurements, respectively. Following the analysis protocols of quasi-paired cohort (see below), a new paired cohort was constructed where individuals from ASD and control groups were cross paired to identify ASD-associated microbiome features (see below and Fig. 1). Microbiome features, such as abundance of metabolic pathways, were compared between samples in each pair, and those significantly overrepresented or deficient in ASD samples were suggested as ASD-associated and possibly involved in ASD pathogenesis. Urine biomarkers and clinical rating scores were used to confirm the role of the identified ASD-associated microbiome features. Finally, the identified features were used to construct a diagnostic model and its performance was evaluated with ROC analysis.

**Fig. 1.**
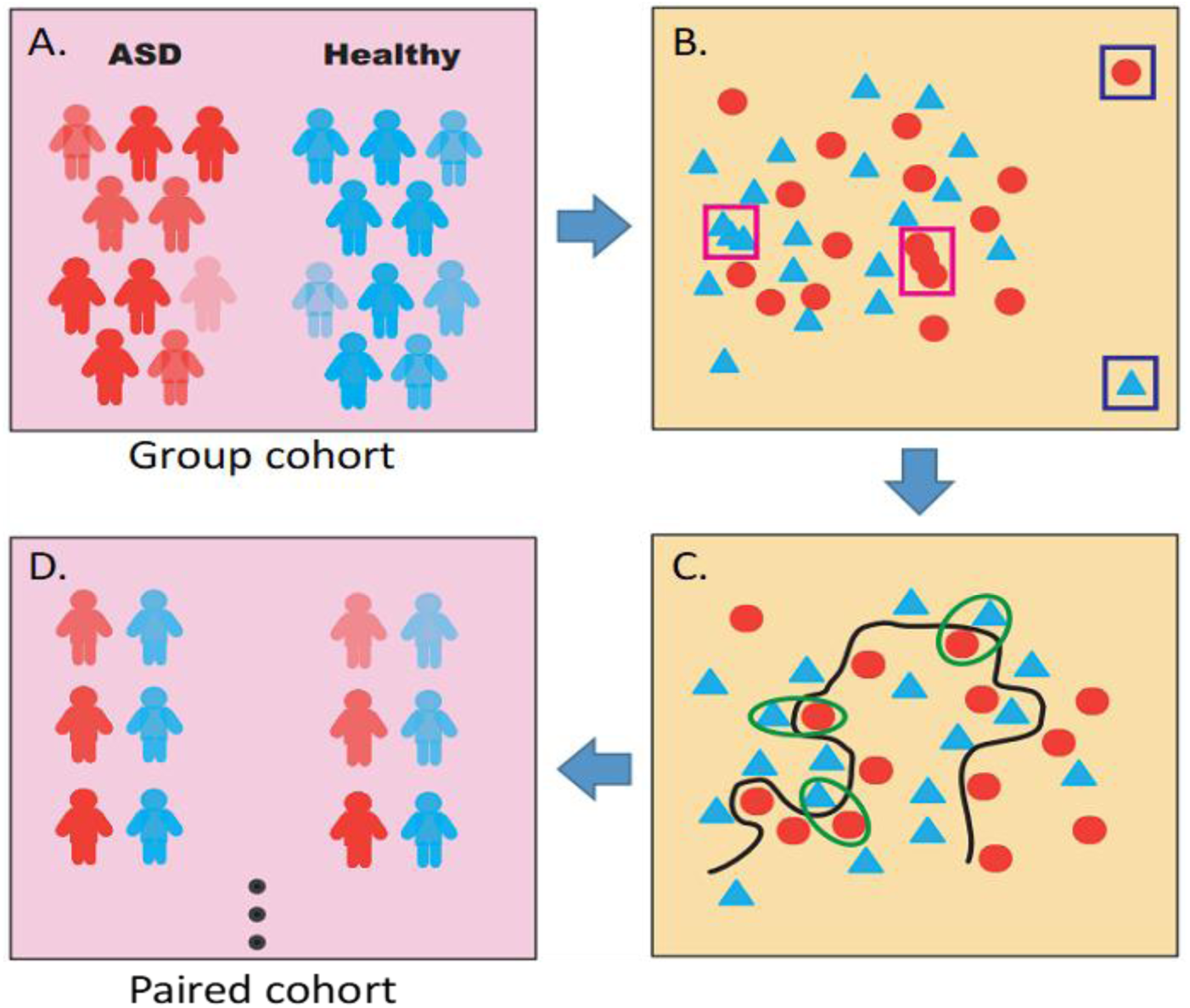
Principle of quasi-paired cohort analysis. **(A)** The group cohort of ASD individuals and healthy control participants. **(B)** High dimensional space with each microbiome feature representing a dimension was constructed and samples were positioned. Samples with abnormally far distances to others (outliners, labeled with blue rectangles) or that were too close (redundancies, labeled with purple rectangles) were removed. **(C)** Boundary samples near the decision boundary (the black line) were selected for both groups and paired with their nearest samples from the opposite group. **(D)** Quasi-paired cohort was constructed for subsequent statistical analysis.

### Participants

Children with ASD and typically developing matched control subjects from 3-8 years old were recruited. The diagnoses of ASD were confirmed with the Autism Diagnostic Interview–Revised ^(*21*)^and Autism Diagnostic Observation Schedule (ADOSTM)(*22*) according to DSM-5 (Diagnostic and Statistical Manual of Mental Disorders, 5th Edition)(*23*) criteria. Rating or scoring instruments of Childhood Autism Rating Scale (CARS)^(*24*)^, Repetitive Behavior Scale-Revised (RBS-R)^(*25*)^, Autism Behavior Checklist (ABC)^(*26*)^, and Gastrointestinal Severity Index (GSI)^(*27*)^ were also performed for autistic children. For the control subjects, medical examinations and parent interviews were performed to exclude anyone with psychiatric disorders. Subjects were also excluded if they experienced infection or took psychoactive medications, antibiotics, probiotics, prebiotics, or a special diet (such as ketogenic diet) in the three months prior to enrollment.

### Metagenome sequencing and annotation

Fecal specimens of all participants were collected and immediately frozen at −20°C in sample tubes and stored at −80°C for subsequent metagenome sequencing. Total DNA of each fecal sample was extracted using the QIAamp PowerFecal Pro DNA Kit (Qiagen) and paired-end sequencing was performed on Illumina HiSeq X10 platform (150 bp×2). Raw reads were first applied to quality control where ambiguous sequences and adapters were filtered out by FastQC(Version 0.11.5) (http://www.bioinformatics.babraham.ac.uk/projects/fastqc/), and low-quality bases and reads were trimmed by the FASTX-Toolkit (Version 0.0.13) (http://hannonlab.cshl.edu/fastx_toolkit/). Then, reads of human origin that mapped to Hs37 human genomes by BWA (mem module with default parameters) (http://bio-bwa.sourceforge.net/) were removed, and PCR duplicates were removed by PRINSEQ (http://prinseq.sourceforge.net/). The final clean reads were applied to taxonomy and metabolic function annotation. For each sample, MetaPhlAn2 (http://www.huttenhower.org/metaphlan2) was used to perform microbial classification and calculate relative abundance of each species, and HUMAnN2 (http://huttenhower.sph.harvard.edu/humann2) was used to annotate microbial pathways according to the BioCyc Database (https://biocyc.org/) and determine the abundance of each pathway. Alpha-diversity of each sample indicated by Shannon index was calculated using the R vegan package (https://cran.r-project.org/web/packages/vegan/index.html), and principal component analysis (PCA) was used to evaluate the beta-diversity among samples.

### Construction of quasi-paired cohort

To construct the “paired cohort” from the original group cohort, we first constructed a high-dimensional space where each microbiome feature (abundance of species or metabolic pathway) represented a dimension. Then all subjects were positioned in the space according to their microbiome constitution profiles, and similarity of each sample to its nearest k neighbors was presented as KNN (the average Bray-Curtis distance to k nearest neighbors). Here, k was the square root of sample size. First, outliers that were too far (KNN>mean KNN+SD) and redundancies that were too near (KNN<mean KNN-SD) to their neighbors were removed to avoid stochastic impacts from these samples on selection of paired sample and statistics. Next, for both ASD and control groups, we identified boundary samples using samples that were more similar to neighbors in the opposite group than their own group, i.e. intragroup KNN > intergroup KNN. Theoretically, the samples located near the boundary between phenotypes (here ASD and control) in the high-dimensional space are very valuable in describing the boundary with microbiome features as they often contain the most deviated features that could clearly divide the two phenotypes. Comparable numbers of boundary samples were selected for each group, and ASD-control pairs were constructed with one boundary sample and one of its k nearest neighbors of the opposite side, thus the pairs are composed of samples of similar microbiome constitutions (nearest neighbors in the high-dimension space) but in opposite phenotype groups. Finally, these ASD-control pairs constituted the quasi-paired cohort after removal of redundancy (Fig. 1, Fig. S1).

### Measurement of urine organic acid

Morning urine of ASD and control children was collected and immediately frozen at −20°C then transferred to a −80°C freezer. For each urine sample, a total of 75 metabolites was measured utilizing Agilent 7890A gas chromatograph–mass spectrometer (Agilent Technologies, Santa Clara, CA, USA) according to the manufacturer’s instructions, and analyzed with MSD ChemStation (E.02.02.1431) to calculate the concentration of each metabolite normalized by the concentration of creatine in the same sample.

### Random forest model

The random forest model was constructed with the caret (https://cran.r-project.org/web/packages/caret/) and randomForest (https://cran.r-project.org/web/packages/randomForest/index.html) R packages to select the most deviated markers of enzymes. The model was trained by 50% of samples through two-fold cross validation, and tested with all samples. Bootstrapping was done 1000 times. For each bootstrap result, the contribution of each enzyme to the model and the diagnostic score for each subject were recorded. Then the diagnostic capacity of the panel of detoxification enzymes we identified was evaluated with AUC (area under ROC curve) and accuracy through the R packages pROC (https://cran.r-project.org/web/packages/pROC/index.html) and ROCR (https://cran.r-project.org/web/packages/ROCR/index.html) in discriminating the ASD patients from control subjects. Mean contributions of each enzyme to the model were calculated to evaluate their deviations between ASD and control, and mean diagnostic score was utilized to comprehensively represent the extent of dysfunction in microbial detoxification as the score was inferred from the abundance of detoxification enzymes and adjusted by the model according to their deviations between ASD and control.

### Statistical Analysis

Wilcoxon rank-sum test was used to compare the average between ASD and control group in their alpha-diversity indicators and abundance of each species, with adjusted *p* <0.05 as significant. Statistical significance in the abundance of pathways between paired samples of the quasi-paired cohort was tested with the Wilcoxon signed-rank test for paired samples with *p* < 0.05 and fdr <0.1. Correlations of pathways vs. urine organic acids, and score in detoxifying dysfunction vs. ASD rating scores were evaluated with Spearman’s rank correlation coefficient, and those with **ρ**<-0.4 or >0.4 were regarded as strong correlations.

## Results

### Intestinal microbiomes showed great individual diversity

The study enrolled 79 participants including 39 children with ASD and 40 age- and gender-matched neurotypical control subjects (mean age, 5.59 years; range, 3-8 years; male %, 82%; Table S1). Metagenome sequencing and analysis of stool samples identified a total of 209 species in all samples with each sample containing an average of 101±14 species. The alpha diversity or richness of the microbiomes were similar between ASD and HC (Fig. S2). Among the 209 species, 18 showed significant differences between groups (Fig. S3), including *Veillonella parvula* and *Lactobacillus rhamnosus* enriched in ASD, while *Bifidobacterium longum* and *Prevotella copri* were enriched in HC (Wilcoxon rank-sum test, p<0.05, fdr<0.3). These findings are partially consistent with previous studies on 16S rDNA sequencing-based species profiling(*8, 28, 29*). However, each of these studies, including ours, identified a list of study-specific differential species for ASD, and these species do not provide clear mechanisms for understanding ASD pathogenesis.

Principal component analysis (PCA) analysis of species profiles did not obviously separate samples of ASD from control group (Fig. S4A) when plotted with the top two PCs, and the samples of each group were widely scattered, indicating great individual diversity even among samples of the same group. We next calculated the pairwise Bray-Curtis distances between each pair of samples and found that intergroup distances were not larger than intragroup differences (Fig. S4B), implying that the individual diversity was so great that it overwhelmed ASD-associated alterations. After functional annotation of the metabolic pathways present in the microbiomes, we inferred the abundance of each pathway for each sample. PCA analysis of the metabolic profiles did not show clusters of samples of the same group (Fig. S4C), and the pathway-based Bray-Curtis distances of paired samples were similar between intergroup and intragroup (Fig. S4D), suggesting great metabolic diversity among samples as well.

### The quasi-paired cohort strategy identified deficient microbial detoxification in ASD

According to our quasi-paired cohort strategy, we constructed a cohort of 65 ASD-control pairs, which ultimately included 20 ASD subjects and 18 control subjects from the original groups based on the metabolic profiles of samples. Comparison between the paired samples identified a total of 96 ASD-associated pathways (Wilcoxon signed-rank test, P<0.05, fdr<0.1, Table S2) with 39 over-represented and 57 deficient pathways involved in many metabolic categories (Table S3).

Among the list of the ASD-associated pathways we identified, a conspicuous trend was the deficiencies in the metabolic category of detoxication in ASD samples. A total of five complete pathways in this category exhibited obviously decreased abundance in ASD subjects when compared to their control counterparts (Fig. 2). Two of the impaired detoxication pathways in ASD were involved in the generation of glutathione (GSH): pathways of the γ-glutamyl cycle and biosynthesis of L-glutamate and L-glutamine, the precursor of GSH (Fig. S5). The three other pathways functioned in degradation of organic toxicants of methylphosphonate, 3-phenylpropanoate or 3-(3-hydroxyphenyl)-propanoate, and methylglyoxal (Fig. S6). Almost all the enzymes in the pathways exhibit significant deficiency in ASD samples including key enzymes in these pathways (Wilcoxon signed-rank test, *p*<0.05): glutamate-cysteine ligase (*gshA*), glutathione synthase (*gshB*), *γ*-glutamyltransferase (ggt), and aminopeptidase B (*pepB*) in GSH biosynthesis; C-P lyase (*phnJ*), which removes the phosphate group in degrading methylphosphonate; dehydrogenase (*hcaB*) and oxidase (*mhpB*), which break the benzene ring; as well as glyoxalase (*gloA*) and hydrolase (*gloB/gloC*), which degrade the methylglyoxal to lactate (Fig. 2).

**Fig. 2.**
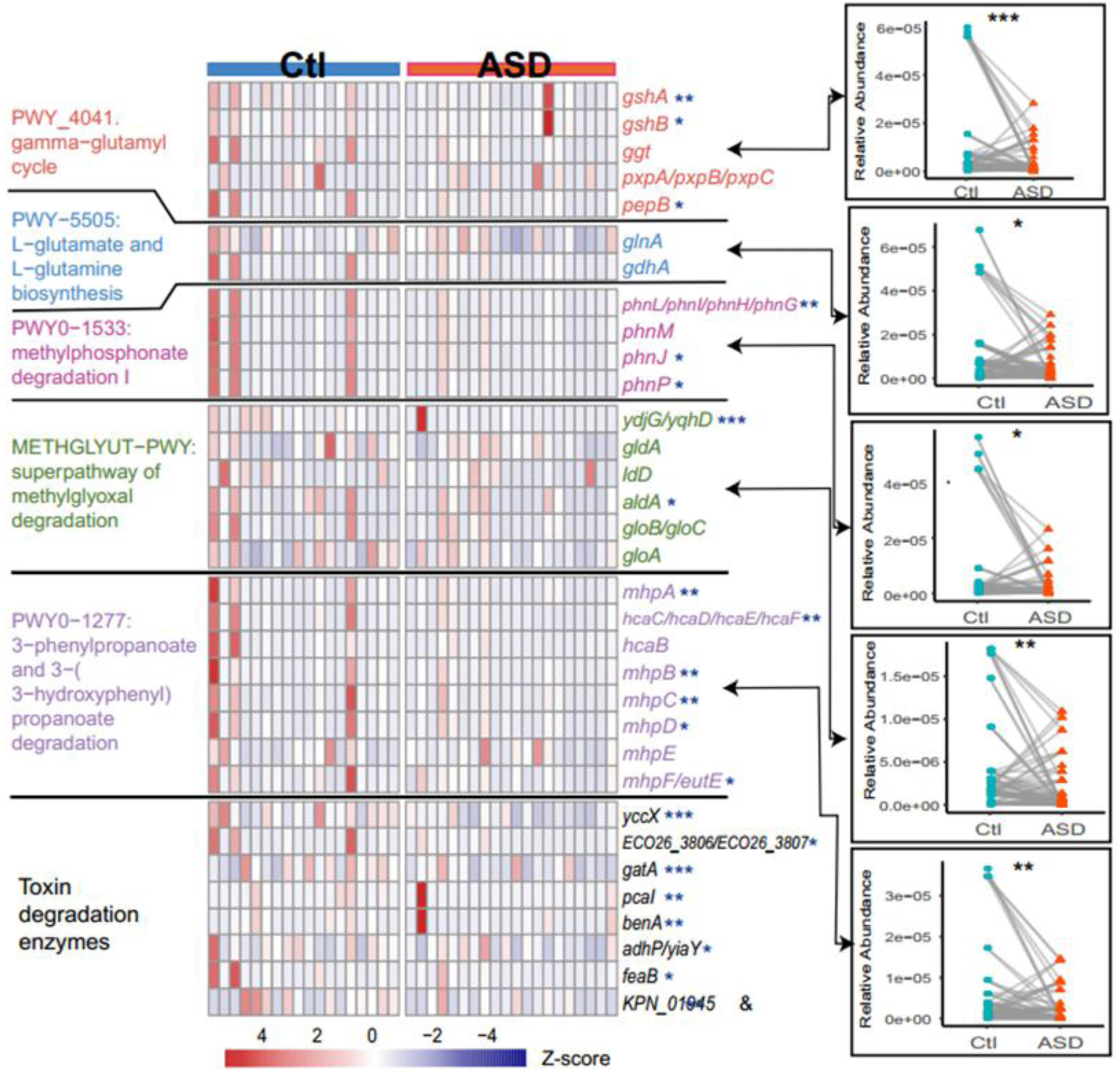
ASD-associated detoxifying enzymes and pathways. The heatmap shows the relative abundance of each detoxifying enzyme in each sample involved in the paired cohort. Colors indicate the relative abundance that is normalized with Z-score. Names of pathway modules and representative genes in Enterobacteriaceae of the enzyme are labeled on left and right sides of the heatmap, respectively. Comparison of the relative abundance of pathways between controls and ASD are shown on the right of the heatmap and linked to corresponding pathways in the heatmap with arrows. Significance was tested with Wilcoxon signed-rank test; *, *p*< 0.05; **; *p* < 0.01; ***, *p* < 0.001.

Apart from the above detoxification pathways, we further scrutinized all enzymes participating in degradation of various kinds of toxins, and compared their abundance between subjects of ASD and their control counterparts (Wilcoxon signed-rank test, *p*<0.05). None of the enzymes exhibit significant overrepresentation, while eight were significantly deficient in ASD, and participate in the degradation of a wide range of toxicants including chloroalkane/chloroalkene, aminobenzoate, benzamide, styrene, naphthalene, xylene, and benzoate (Fig. 2, Table S4). These toxicants are widely used as insecticides and food additives. The deficiency in these critical enzymes suggests a wider range of impairment in detoxification in ASD, although their relevant pathways were not significantly different between twin samples.

The other ASD-associated pathways we identified were largely consistent with previous knowledge about ASD. Many of these pathways were involved in the biosynthesis of pyrimidine, purine, and tetrahydrofolate (Table S3) that were deficient in ASD children. It is well-known that metabolism of tetrahydrofolate is often defective in ASD patients(*30*). Tetrahydrofolate is the one-carbon donor for the biosynthesis of nucleotides and homocysteine re-methylation to form methionine and subsequently S-adenosylmethionine (SAM), and a major source of folate—the precursor of tetrahydrofolate is food, which confirms the deficiency in microbial metabolism of tetrahydrofolate in addition to host defect in ASD. Among the 39 ASD-enriched pathways, 17 are from yeast and one of them synthesizes 2-amino-3-carboxymuconate semialdehyde (Table S3), the intermediate to generate an excitotoxin—quinolinic acid—through the subsequent kynurenine pathway. This result is consistent with the previous findings in ASD patients of overproduction of quinolinic acid(*31*) and overgrowth of yeast in the intestine(*32*). The consistency of the ASD-associated pathways we identified with previously reported metabolic alterations in ASD supported the reliability of our novel analysis strategy of quasi-paired cohort, and further confirmed the contributions of gut microbial metabolism to hosts.

### Impaired microbial detoxification is associated with mitochondrial damage and the extent of ASD severity

Metabolism is obviously disturbed in ASD, and we measured the absolute concentrations of urine metabolites for our subjects to quantitatively evaluate their metabolic alterations. The results showed that most ASD children exhibited metabolic abnormity when compared with controls, which is in accordance with previous findings(*33*) (Table S5). The most significantly abnormal metabolites in our ASD subjects included aconitic acid, suberic acid, 2-hydroxyhippuric acid and fumaric acid, all biomarkers of damage or dysfunction in mitochondria. Of importance, the quantities of these biomarkers showed significant negative correlations with the abundance of most of the ASD-associated detoxication enzymes we identified (Fig. 3). The correlations suggested a potential role of these microbial detoxification enzymes for protecting mitochondria from oxidative and toxic stresses, and the loss of such protection in ASD may contribute to mitochondrial damage, one of the major pathological alterations in various tissue of ASD children, including the brain.

**Fig. 3.**
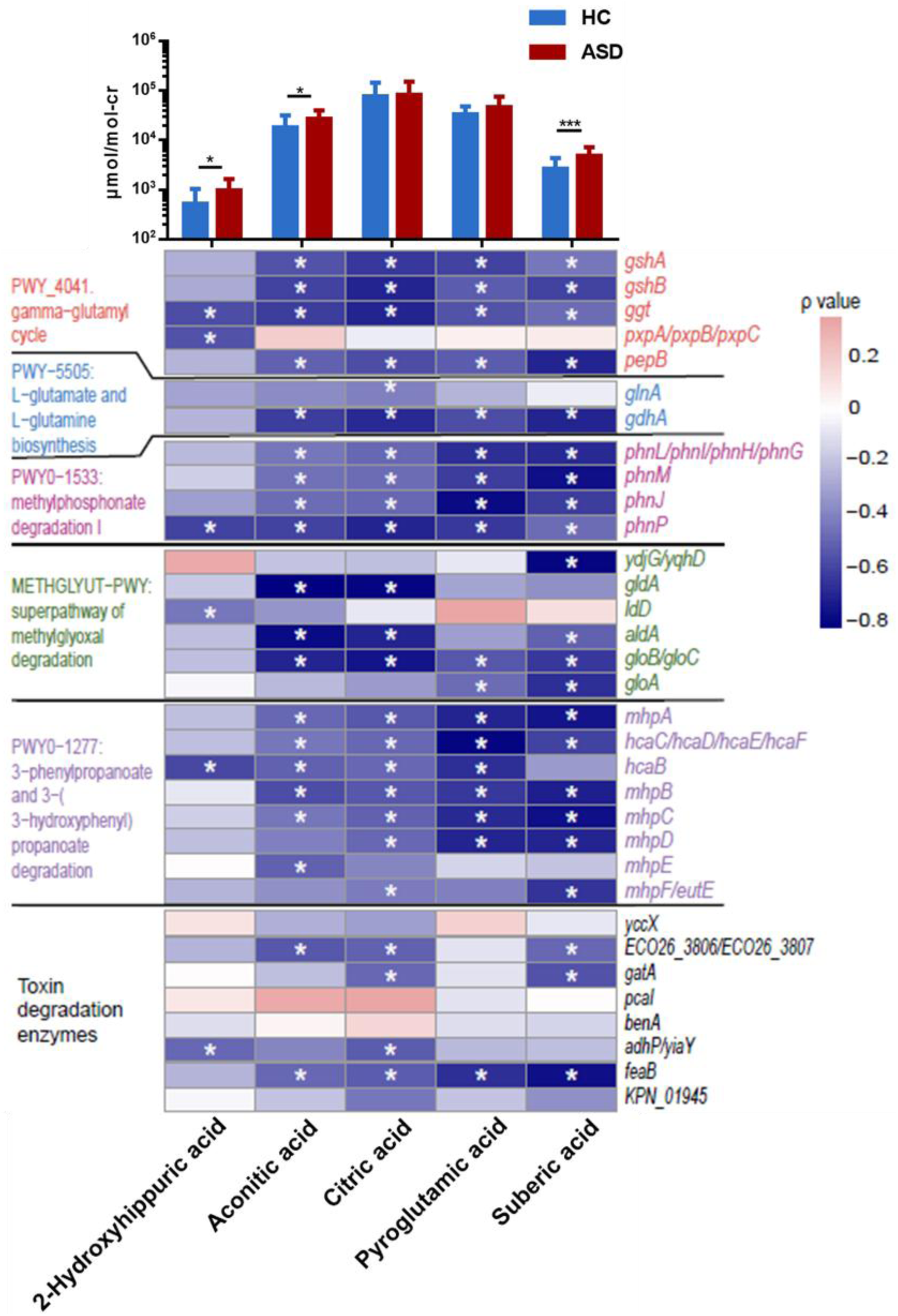
The correlation of detoxifying enzymes to urine biomarkers of mitochondrial damage. The upper panel shows the average concentrations of each urine biomarker (labeled below the heatmap) in ASD and control samples. The y-axis indicates the concentrations (μmol normalized by the concentration of creatine of the same sample). Significance was labeled with *, *p*< 0.05; **; *p* < 0.01; ***, *p* < 0.001 based on a Wilcoxon signed-sum test. The lower panel is the heatmap of correlations between each detoxification enzyme (in the same order as Fig. 2) and biomarkers of mitochondrial damage. The color indicates the ρ value of the Spearman rank test, *, ρ≥ 0.4 or ρ≤ −0.4.

To corroborate these detoxification enzymes to the development of ASD, we constructed a random forest classifier based on the abundance of the detoxification enzymes we identified (as listed in Fig. 2) and evaluated its performance in discriminating ASD from control subjects. The ROC evaluation with 1000 bootstrap replicates showed that the panel of detoxification enzymes accurately described the deviations between ASD and control subjects, and achieved a diagnostic power as high as 88% AUC (Fig. 4A). The diagnostic model also outputs the contributions of each enzyme to the model with the top five as essential enzymes in biosynthesis of GSH and L-glutamate/L-glutamine, and degradation of aminobenzoate, chloroalkane/chloroalkene/naphthalene, and methylglyoxal, which means these enzymes are most deviated between ASD and controls (Fig. 4B).

**Fig. 4.**
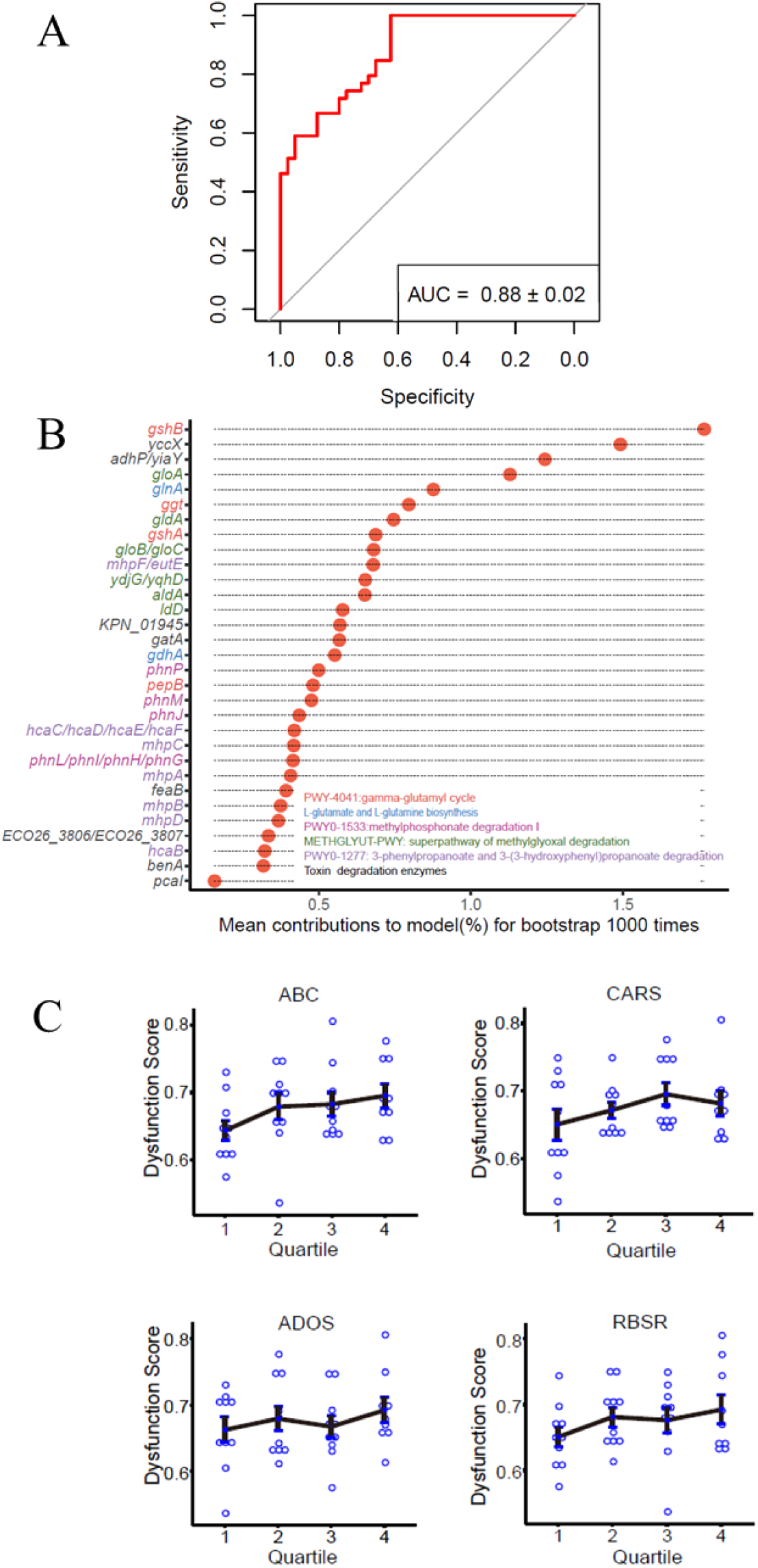
Diagnostic model of detoxifying enzymes for ASD status. All identified detoxifying enzymes were used to construct a diagnostic model for prediction of ASD status by random-forest classifiers with 1000 bootstrap replicates. **(A)** The ROC analysis of the performance of the diagnostic model. AUC, area under the curve. **(B)** The mean contributions of each detoxifying enzyme based on 1000 bootstrap replicates. **(C)** Dysfunction scores of the microbial detoxification inferred from the diagnostic model for samples in each quartile of patients divided according to their clinical ASD rating scores.

Further, we investigated the correlations between the mean diagnostic score of each sample in the model and the severity of ASD. As the diagnostic score was inferred from the abundance of detoxification enzymes and adjusted by their respective deviation between ASD and control, the score can be regarded as a comprehensive score for the extent of impairment in microbial detoxification, and is hereinafter named the dysfunction score. As there are no objective biomarkers for the disease severity or even diagnosis of ASD, we applied clinical rating scores for ASD, i.e. ADOS, ABC, CARS, and RBSR, to divide patients into respective quartiles according to each of the scores. The dysfunction scores of patients in quartiles showed a trend of increasing with the rating scores (Fig. 4C). Logistic regression between the dysfunction score and clinical rating scores largely confirmed their positive correlations, although they were not significant possibly due to the subjectivity in clinical rating of ASD (Fig. S7). These correlations between impaired microbial detoxification and the extent of severity of ASD further demonstrated the contribution of the intestinal microbiome in the pathogenesis of ASD through the dysfunction of detoxification.

## Discussion

Our study, supported by the power of the quasi-paired cohort strategy, revealed a previously unidentified ASD-associated deficiency in microbial detoxification, which exhibited a strong correlation to the extent of mitochondrial damage, as well as the severity of clinical ASD manifestations. Toxicants exposure has been epidemiologically confirmed as an important etiological factor of ASD and patients often show clinical manifestation of intoxication^53^. In fact, mammals are consistently exposed to toxicants from the external environment like glyphosate, or from internal metabolic process of host or microbes, such as methylglyoxal. Detoxication function is thus essential for life. In addition to the host detoxication system such as hepatic enzymes like P450, the intestinal microbiome may function as the first frontier of degrading or expelling toxins as the digestive tract is the major route by which we uptake toxins.

The mitochondria are a major target of organic toxicants due to their lipophilic property and depletion of GSH, and damaged mitochondria will release various organic acids into circulation, which are discharged from urine. Our finding of impaired microbial detoxification provides an explanation of why ASD children are so vulnerable to environmental toxins and suggest microbial detoxication is a considerable part of the entire host detoxification system and contributes to the pathogenesis of ASD if it is severely impaired (Fig. 5). However, the reasons underlying the specific dysfunction in microbial detoxification is not clear, and might be one of the consequences of microbiome dysbiosis caused by various genetic and environmental factors such as altered diet and defects in digestion, which change the nutrients provided to the microbial inhabitants.

**Fig. 5.**
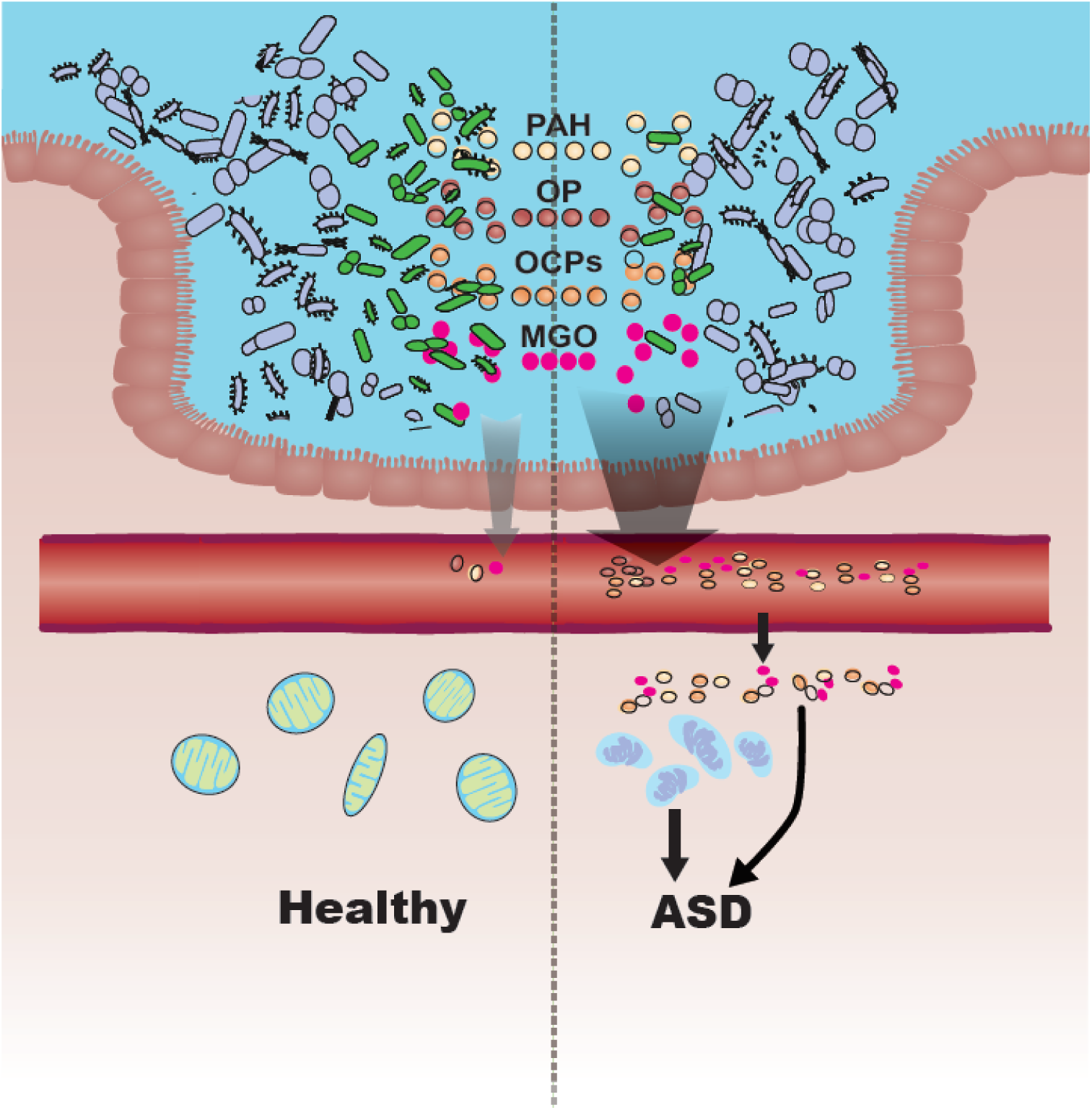
Mechanism of how impaired microbial detoxification contributes to the pathogenesis of ASD. Colored dots in the intestinal lumen indicate common toxicants indigested. PAH, polycyclic aromatic hydrocarbon; OP, organophosphorus; OCPs, organochlorine pesticides; and MGO, methylglyoxal. Intestinal bacteria in green indicates microbes able to detoxify the toxicants, which are obviously deficient in ASD. More toxicants are thus absorbed into circulation in ASD due to the defect in microbial detoxification. Accumulation of toxicants in tissues cause damages in mitochondria or other structure/function integrity, which finally leads to ASD.

Toxicants implicated in ASD include organochlorine pesticides, polycyclic aromatic hydrocarbons, automotive exhaust, and heavy metals, most of which are organic compounds, especially aromatic and halogenated^54^. Organic toxicants are often lipophilic and prone to accumulate in fatty tissue such as the brain following chronic intake and then damage membrane structures such as those of mitochondria. Glyphosate (N-phosphonomethyl-glycine), the most-widely used herbicide in the world, is a methylphosphonate well known for its nuclear and mitochondrial toxicity. Microbial degradation of methylphosphonate is very common in bacteria such that many species utilize it as sole source of phosphorous^(*34*)^, but our findings shows that such a pathway is obviously deficient in ASD.

Aromatic hydrocarbons are compounds containing one benzene ring or more than one fused ring (polyaromatic hydrocarbons—PAHs). Many bacterial species especially in the phyla of *Proteobacteria* and *Actinobacteria* are able to break down aromatic hydrocarbons through the pathway of 3-phenylpropanoate and 3-(3-hydroxyphenyl)-propanoate degradation, which is also defective in ASD. Methylglyoxal is an inevitable by-product of the processes of glycolysis and metabolism of fatty acid and protein. Methylglyoxal can permeate cells and is deleterious as a highly potent glycating agent reacting with protein, lipids, and nucleic acids to form advanced glycation end products (AGEs) which are implicated in various degenerative processes including lesions in the brain^(*35*)^. One study found that methylglyoxal changes the function and oxidative state of mitochondria and causes damage to mitochondria in the brain of ASD individuals ^(*36*)^. Therefore, gut microbes are essential to quickly degrade the local methylglyoxal *in situ* to keep the concentration of methylglyoxal at a low non-toxic level.

Apart from direct toxin degradation, the ASD microbiome is also deficient in the biosynthesis of GSH. GSH is one of the body’s major antioxidants and is the key cofactor for a number of detoxifying enzymes. GSH is essential in degrading organic toxicants and expelling heavy metals, both of which contribute to maintaining the function of mitochondria^(*37*)^. As the precursor of GSH, L-glutamine is not only involved in organic toxin degradation but is also beneficial to leaky gut and ulcers. Glutamine has the ability to maintain intestinal barrier integrity and function to stop organic toxicant such as LPS (lipopolysaccharides) into circulation^(*38, 39*)^. Thus, the GSH generated by intestinal microbes adds a great contribution to local detoxification.

Growing evidence has suggested the role of mitochondrial dysfunction in the pathogenesis of ASD with both inborn defects in mitochondria and acquired damage caused by environmental toxins reported in cases of ASD ^(*33*)^. Studies have found that damaged mitochondria may release mtDNA and other DAMPs (damage-associated molecular patterns), which activate a low-grade inflammatory response in various tissues including the brain^(*40*)^. Such systemic low-grade inflammation with elevated inflammatory cytokines, chemokines, and cellular immune activity, is well-documented in ASD patients^(*41*)^, and the function of neurocytes and brain development may be thus impaired. Recently, cross-talk between microbiome and the mitochondrion has been reported^(*42*)^ as specific microbial products may inhibit or fine-tune the function of mitochondria^(*43-45*)^. Here, we reported a protective effects of the microbiome on mitochondrial structural and functional integrity, which is illustrated by the close correlation between impaired microbial detoxification and mitochondrial damages.

Although this study has discovered a deficiency in the detoxification pathways of the intestinal microbiome in ASD patients from metagenomic data, evidence of toxicant accumulation is not easy to obtain. First, many toxicants and their reductive antioxidants we identified are very chemically active, such as methylglyoxal and GSH, and they rapidly react with surrounding biomolecules, which makes them hard to detect in the frozen samples. Second, the amount of toxicant exposure often fluctuates, and toxicants are possibly undetectable in feces once absorbed although the chronic damage they cause may last for a long time, which uncouples the toxicant concentration in samples from clinical manifestations of intoxication. Third, the substances of the detoxification enzymes we identified is a long list of organic toxicants and exposure to them varies from person to person, which makes the study design of any transection cohort difficult for quantitative evaluation of the long term exposure to various toxicants and its correlations to chronic damage and to clinical manifestations of ASD. Finally, no suitable method is currently available to simultaneously measure the absolute concentrations of the metabolites we are interested in. Thus, to clarify the pathogenesis of ASD in the aspect of toxicant exposure and detoxification, novel methods of toxicant measurement are needed, which would be useful not only in demonstrating toxicants accumulation but also valuable for clinical evaluation of individual etiological factors as well.

In conclusion, our study proposed a new strategy for metagenome analysis—quasi-paired cohort, and successfully identified a conspicuous trend of impairment in detoxification in the ASD gut microbiome. The impaired microbial detoxification correlated to clinical rating of ASD and extent of mitochondrial damage—one of the main pathological alterations of ASD, which strongly suggests that impaired microbial detoxification contributes to the pathogenesis of ASD. Such a previously unknown mechanism points to potential future therapeutic strategies of rebuilding the impaired microbial detoxification for ASD patients. Moreover, the novel strategy of quasi-paired cohort is powerful for metagenome analysis, and might be applied to studies of other complex diseases or even other -omics datasets with similar characteristics of high-dimensionality and complexity.

## Supplementary Materials

Fig. S.1 Flowchart of quasi-paired cohort analysis

Fig. S2. Diversity and richness of the microbiome of ASD and HC. Fig. S3. Differential species in HC and ASD samples.

Fig. S4. Principal component analysis (PCA) analysis of species and metabolic profile.

Fig. S5. Metabolic pathway of γ-glutamyl cycle (A) and biosynthesis of L-glutamate and L-glutamine (B).

Fig. S6. Metabolic pathway of degradation of methylphosphonate (A), 3-phenylpropanoate or 3-(3-hydroxyphenyl)-propanoate (B), and methylglyoxal (C).

Fig. S7. Logistic regression between the dysfunction score and clinical rating scores.

Table S1. Characteristics of participants.

Table S2. ASD-deficient and ASD-associated pathways in Metabolic Twins

Table S3. The metabolic categories which ASD-associated pathways involved in.

Table S4. Enzymes deficient in ASD involved in degradation of toxicants.

Table S5. Profile of urine organic acid test of ASD children.

## General

This study has been approved by the Institutional Review Board of Peking University (Ethical Review Document No: IRB00001052-17100). The study group gained informed consent from the parents/guardians for the collection of stool and urine samples and trial information. We confirmed that all methods were performed in accordance with relevant guidelines and regulations. The authors are grateful to all the children who participated in this study and their parents for their cooperation.

## Author contributions

JW and YK conceived designed the project. MZ and RD collected information from the participants. YC, QM and XS processed and analyzed the whole metagenome sequencing data. YC and XS performed the statistical analysis. YK and JY conceived and coordinated the project. ZW recruited, diagnosed and examined the recruited participants. All authors contributed to the writing of the manuscript.

## Data and materials availability

The raw metagenome sequencing data reported in this paper have been deposited in the Genome Sequence Archive in BIG Data Center, Beijing Institute of Genomics (BIG), Chinese Academy of Sciences, under accession numbers CRA001746 at http://bigd.big.ac.cn/gsa.

